# Satellite Cell Expression of RAGE is Important for Collateral Vessel Formation

**DOI:** 10.1101/2021.03.02.433612

**Authors:** Laura Hansen, Giji Joseph, W. Robert Taylor

**Affiliations:** Division of Cardiology, Department of Medicine, Emory University, Atlanta, GA; Division of Cardiology, Atlanta Veterans Affairs Medical Center, Decatur, GA; The Wallace H. Coulter Department of Biomedical Engineering, Georgia Institute of Technology and Emory University, Atlanta, GA

**Keywords:** RAGE, Satellite Cells, Collateral Vessels

## Abstract

**Objective:** The growth and remodeling of vascular networks is an important component of the prognosis for patients with peripheral artery disease. One protein that has been previously implicated to play a role in this process is the receptor for advanced glycation end products or RAGE. This study sought to determine the cellular source of RAGE in the ischemic hindlimb and the role of RAGE signaling in this cell type.

**Approach and Results:** Using a hind limb ischemia model of vascular growth, this study found skeletal muscle satellite cells to be a novel major cellular source of RAGE in ischemic tissue by both staining and cellular sorting. While wild type satellite cells increased TNFα and MCP-1 production in response to ischemia *in vivo* and a RAGE ligand *in vitro*, satellite cells from RAGE knockout mice lacked the increase in cytokine production both in vivo in response to ischemia and in vitro after stimuli with the RAGE ligand HMGB-1. Furthermore, encapsulated wild type satellite cells improved perfusion after hind limb ischemia surgery by both perfusion staining and vessel quantification but RAGE knockout satellite cells provided no improvement over empty capsules.

**Conclusions:** Thus, RAGE expression and signaling in satellite cells is crucial for their response to stimuli and angiogenic and arteriogenic functions.

**Graphical Abstract:** 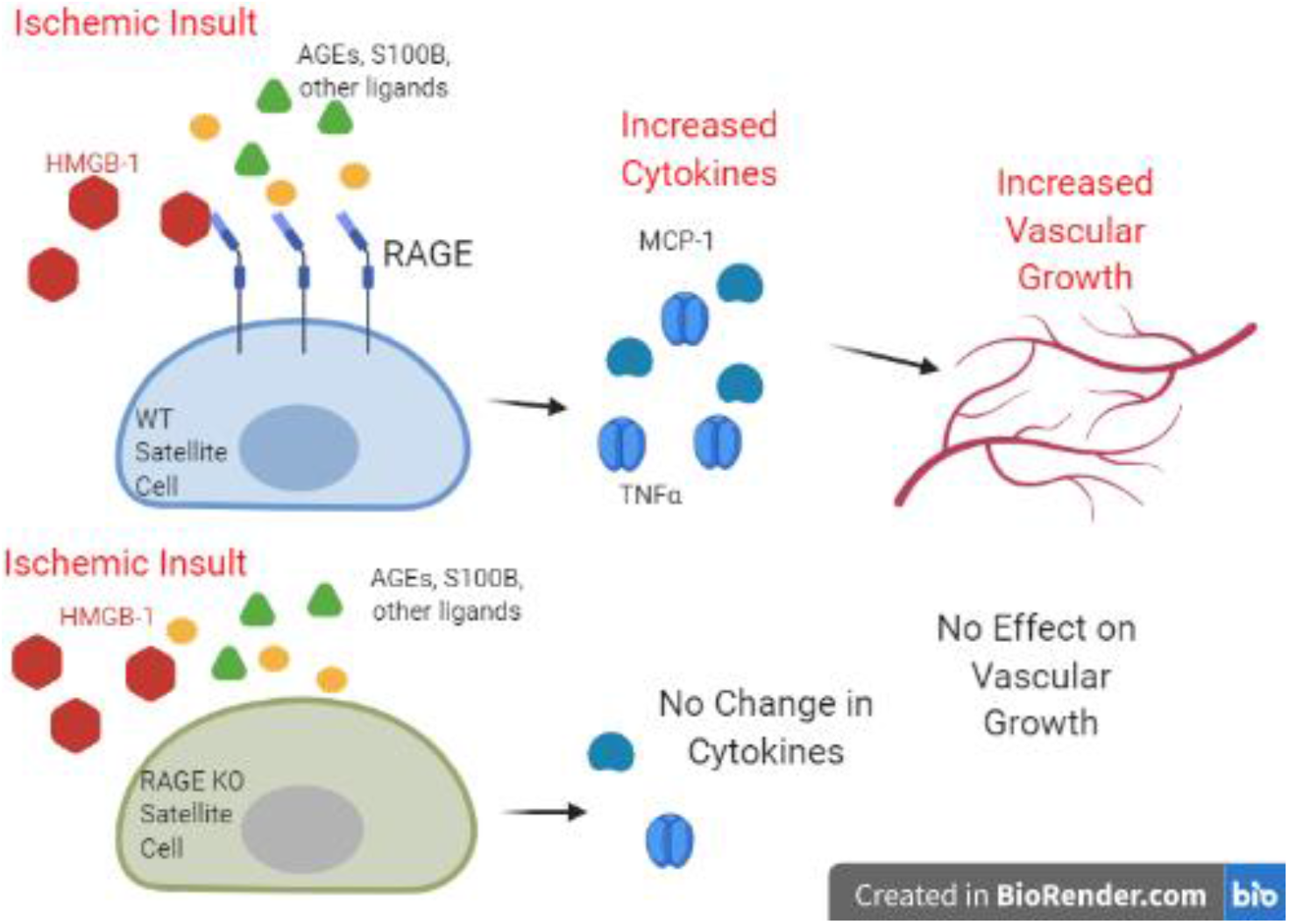

## Introduction

Peripheral artery disease (PAD) is most commonly the result of atherosclerosis in the arteries of the limbs (primarily the legs) impairing blood flow to the extremity or in the carotid arteries impairing flow to the head and brain. Limb PAD often first manifests as pain with physical exertion but chronic oxygen deficiency due to inadequate blood supply can lead to tissue damage that results in ulcers, infections, and amputations. However, the ischemic insult does promote the growth of new blood vessels and the remodeling of existing ones to help restore blood flow and prevent further damage. Indeed, the development of collateral vessels and vascular networks is correlated with better patient outcomes.(1)

The regulation of this vascular response is likely multifactorial with many common cardiovascular risk factors such as smoking, hypertension, hypercholestemia, and diabetes playing a negative role in vessel growth and development. (2, 3) One protein that has been thought to play a role in the poor outcomes observed in diabetic patients is the receptor for advanced glycation end products (RAGE). RAGE is a member of the immunoglobulin superfamily of receptors and has several ligands including s100B, HMGB1 (high mobility group box 1) and advanced glycation end products that are often studied in the context of diabetes.(4, 5) Several studies have investigated the importance of RAGE in vascular injury responses in diabetic animals. RAGE activation and signaling has been shown to result in endothelial dysfunction in coronary arterioles in diabetic mice.(6) RAGE signaling has also been shown to be associated with neointimal hyperplasia and treatment with sRAGE (the soluble form of the receptor) decreases the medial thickening in both diabetic and non-diabetic rats. (7) In an ischemia reperfusion model of myocardial infarctions, both RAGE knockout mice or RAGE inhibition by treatment with sRAGE showed decreased ischemic damage and improved myocardial function compared to wildtype or untreated animals.(8) Specifically related to angiogenesis and vessel growth, the blockage of AGE formation with aminoguanidine improved reperfusion in a femoral artery ligation model. (9) Diabetic RAGE knockout mice or mice that overexpress sRAGE have increased angiogenesis in matrigel plugs with compare to wildtype mice with diabetes.(6) More recent studies have also shown that RAGE knockout mice have improved outcomes following hind limb ischemia procedures.(10, 11) One interesting finding in these recent studies was RAGE seemed to play a role in even in the non-diabetic mice likely due to signaling from increases in the RAGE ligand HMGB1.(10) Expression of RAGE in cells such as endothelial, smooth muscle, and macrophages is currently thought to be the major player in RAGE signaling; however, we have identified a novel cell type with significant RAGE expression: satellite cells. The current study investigates unique role of RAGE signaling in skeletal muscle satellite cells and the paradoxical, proangiogenic effects.

Satellite cells are skeletal muscle stem cells which play an important role in the repair of muscle tissue. Healthy muscle contains a quiescent pool of satellite cells that are capable of repairing by the muscle by proliferating and differentiating into myotubes in response to injury as well as self-renewing to maintain their population. Yet, the importance of RAGE signaling by these cells in relation to vascular growth and repair has not been studied. The objective of this study was to investigate the role of RAGE signaling and vessel formation in a non-diabetic setting. We identified skeletal muscle satellite cells as a novel cellular site of RAGE expression in a hind limb ischemia model of vessel formation and studied the function of RAGE expression in skeletal muscle satellite cells.

## Methods

### Surgical Model

All animal studies were conducted under the approval of the Emory University Institutional Animal Care and Use Committee. Mice were either bred in house (RAGE KO) or obtained from Jackson Laboratories (C57Bl/6) or Charles River (129Sv2). Male mice between 8 and 10 weeks of age were anesthetized with 1-2% isoflurane via inhalation in a chamber and then maintained through a nose cone during the procedure. The animals received Buprenex (0.1mg/kg, subcutaneously) pre-operatively for analgesia and aseptic techniques were employed. The hind limb ischemia procedure was performed unilaterally and the other limb was used as the contralateral control. An incision was made over the left thigh of the mouse exposing the superficial femoral artery and vein. Ligations were made with 6-0 silk suture proximal to the deep femoral artery branch point and just proximal to the branching of the tibial arteries. Following ligation, the length of the artery and vein between the two points was excised. The skin was closed with monofilament nylon suture.

### Satellite Cell Isolation and Culture

Satellite cells for both the establishment of primary cell cultures and for protein and gene analysis were isolated from the adductor and grastrocnemius muscle (the two major muscles affected by the HLI model). The muscles were excised and digested using both mechanical and enzymatic methods.(12) Muscles were first minced before digestion in 0.1% Pronase (Calbiochem). The digested cells were dissociated and passed through a 100 μm filter. Satellite cells were purified from this cell suspension using magnetic bead separation (Satellite Cell separation kit from Miltenyi Biotec). For protein and gene expression assays of freshly isolated cells, the cells were pelleted and stored in a −80 °C freezer until use in the assays described below. This process also resulted in viable cells, which were capable of continued growth and proliferation in culture. The primary cell cultures were maintained in Hams/ F-10 media (Hyclone) with 20% fetal bovine serum (Sigma), penicillin/streptomycin (Hyclone), and HEPES (Hyclone). These cells were then used for cell delivery and in vitro experiments.

### Histology, Immunohistochemistry, and In Situ Hybridization

Mice were euthanized at day 14 and prepared for histology by perfusion with saline followed by 10% buffered formalin for fixation. For histology and immunohistochemistry, the limbs were demineralized in a formic acid-based solution (Cal-Ex II, Fisher Scientific) for 48 hours before paraffin processing and embedding. The tissues were cut in 5 μm sections for staining. Enzymatic antigen retrieval with proteinase K (Biolabs, 2μg/mL) was used prior to incubation with antibodies. Capillaries were visualized by staining with lectin (Biotinylated Griffonia simplifolia Lectin 1; Vector lab), followed by a Streptavidin Qdot 655 (Invitrogen). Arterioles and arteries were stained for smooth muscle α-actin with a mouse monoclonal antibody (Sigma) using the avidin–biotin–alkaline phosphatase method (Vectastain ABC-AP; Vector Laboratories) and a hematoxylin counterstain. For in situ hybridization, the gastrocnemius muscle was isolated following perfusion fixation and fixed for an additional 24 hours in 10% buffered formalin before processing and paraffin embedding. The blocks were cut in 5 μm sections and baked at 60 °C for 24 hours before storage in −20 °C until use. In-situ hybridization imaging of RAGE was conducted using the QuantiGene ViewRNA ISH Tissue 1-plex Assay kit and procedure from Affymetrix. We performed the assay using a modified 1-plex protocol for Fast Blue Detection using Type 1 Mouse AGER probe and Type 1 Mouse GAPDH probe (Affymetrix). The sections were counterstained with Nuclear Fast Red. A consecutive section was used to stain for satellite cells with a PAX7 antibody (Sigma- Pax7 AB2) and a biotinylated secondary antibody (Vector Labs) and Streptavidin Qdot 655 (Invitrogen).

### PCR

Gene expression was quantified using qRT-PCR analysis. RNA was isolated using RNeasy (Qiagen) from homogenized adductor and gastrocnemius muscle tissue or satellite cell pellets isolated hind limbs. cDNA was purified with QiaQuick (Qiagen) and expression quantified on an Applied Biosystems thermocycler. All primers were purchased from Qiagen.

### Cytokine Array

A UPlex assay (Meso Scale Diagnostics) was used to quantify the protein levels of a number of cytokines. The array contained GM-CSF, IL-1β, IL-6, TNF-α, VEGF, and MCP-1. The protein lysates for this study were obtained by isolating and sorting satellite cells from either the ischemic or non-ischemic limb at post-operative surgical day 7. The cells were lysed in RIPA buffer containing protease inhibitors and equal amounts of total protein lysate were added to each well of the assay plate. The assay was performed using the standard protocol from Meso Scale Diagnostics.

### HMGB-1 Stimulation In Vitro

Cultured wild type and RAGE KO satellite cells (p5-p8) were plated in collagen coated 6 well plates. Cells were stimulated with 250 nM HMGB-1 (Sigma) for 24 hours before RNA isolation. RNA was isolated as described above using a RNeasy kit (Qiagen) and cDNA was purified with a QiaQuick kit. (Qiagen). Qiagen Quantitect primers were used to quantify expression using an Applied Biosystems thermocycler system.

### Alginate Encapsulation and Cell Delivery

To test the differential effects of satellite cells with and without RAGE on collateral growth and recovery, we delivered encapsulated satellite cells at the time of HLI surgery. We selected this approach as we have previously shown that alginate encapsulation enhances cell survival.(13, 14) Cultured satellite cells were suspended in 1% ultrapure low viscosity alginate (Novamatrix) and collected in a gelling solution of 50 mM BaCl_2_. An electrostatic encapsulator (Nico) with a 0.17 mm nozzle, 10 mL/h flow rate, and 7 kV voltage generated microcapsules approximately 200-250 μm in diameter. Capsules were washed in 0.9% saline solution and stored in saline on ice during the HLI surgery until implantation. To deliver the cells, a small secondary incision medial to the proximal end of the ligation incision was made to create a pocket to deliver 1 million encapsulated cells per animal. Both wild type and RAGE KO primary satellite cell lines generated as described above were used in addition to empty capsules as controls. For this study, the cells were delivered to 129Sv2 mice (Charles River) as this strain has less robust collateral vessel growth thus allowing a better opportunity to observe a therapeutic effect. (15)

### Assessment of Cell Viability

The viability of the cells at the time of encapsulation was verified using a LIVE/DEAD Viability/Cytotoxicity Kit (Invitrogen) and imaged using a Zeiss LSM 510 Laser Scanning Confocal Microscope. Luciferase imaging was used to monitor the viability of cells in vivo. For these experiments, GFP/Luciferase dual expressing satellite cells were derived from transgenic mice from Jackson Labs. Mice received 20 mg/mL of luciferin (GoldBio) via an intraperitoneal injection 45 minutes prior to imaging. Mice were then anesthetized with isoflurane and placed in the In-Vivo Xtreme II whole animal imager (Bruker). Both luminescence and x-ray images were acquired at each time point. Mean luminescence intensity was quantified as a measure of cell viability over time.

### LASER Doppler Perfusion Imaging

Perfusion of the hind limbs was monitored non-invasively using a LASER Doppler perfusion imaging system (Moor Instruments). Mice were anesthetized with isoflurane and placed on a heating pad for 4 minutes. Hair on both hind limbs was removed with depilatory cream and washed with water. Perfusion was assessed as the ratio of the mean perfusion value from equivalent regions of interest (ROI) in the ischemic to non-ischemic limb.

### Statistics

The data was analyzed using t-tests or ANOVA with an appropriate post hoc test (Tukey’s Multiple Comparison Test) to determine significance between individual groups. Data were checked for a normal distribution before statistical analysis. A p-value of less than 0.05 was considered significant and data are reported as mean ± standard error of the mean.

## Results

Hind limb ischemia surgery was performed on mice and the adductor and gastrocnemius muscles were isolated 7 days post-surgery for RNA isolation to quantify RAGE expression. Muscles from the ischemic limb had significantly increased RAGE expression as compared to the muscles isolated from the non-ischemic contralateral limb (p<0.05). (Figure 1A) However, the relatively low expression levels raised the possibility that the source of RAGE was a rare cell type. In situ hybridization of histological slides of the ischemic tissue was used to visualize the RAGE expression in the ischemic tissue. The *in situ* staining showed a pattern similar to that of satellite cells. (Figure 1 B and C). To confirm this observation, satellite cells were isolated from hind limbs following the hind limb ischemia procedure using magnetic bead sorting and analyzed for RAGE expression. Figure 1 D-E shows that the number of satellite cells was increased in the ischemic limb and that the satellite cells isolated from the ischemic limb had increased RAGE expression compared to the non-ischemic limb. Figure 1 F compares RAGE expression in the satellite cell to the non-satellite cell populations from the ischemic limb and found satellite cells had much higher levels of RAGE (p<0.05).

**Figure 1:**
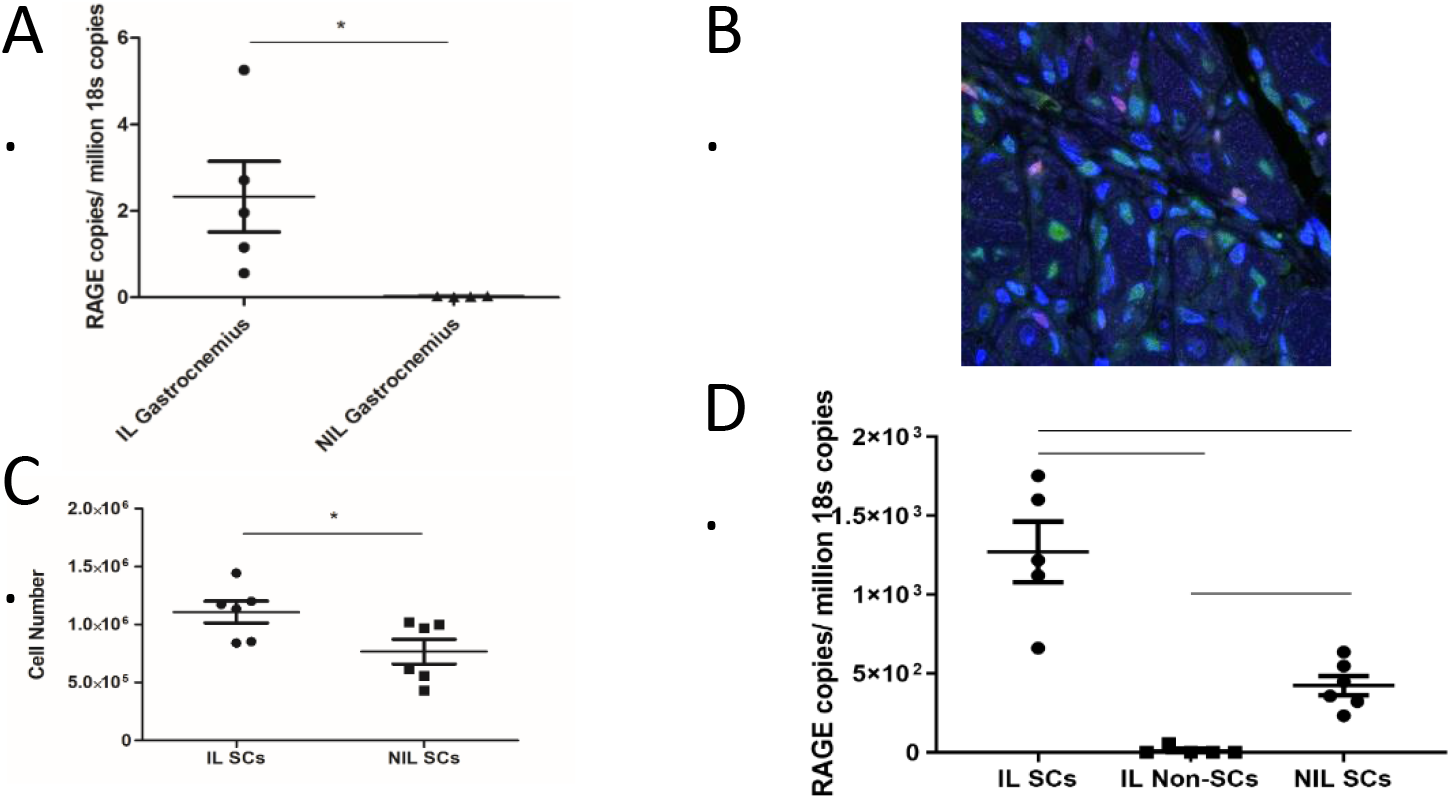
RAGE is Increased in Satellite Cells in the Hindlimb. A) RAGE expression in the hind limb gastrocnemius muscles was determined using qRT-PCR in wild type (WT) mice. The ischemic limb (IL) had increased RAGE expression compared to the non-ischemic limb (NIL). (N = 6, * indicates p<0.05) B &C) Hindlimb muscle was harvested 7 days post HLI and *in situ* hybridization was used to visualize RAGE within the muscle tissue. The representative image shows RAGE (blue dots) and black arrows point to satellite cells with staining. B) Pax7 staining of satellite cells (red) in similar region as RAGE staining (D-F) Satellite cells from the limbs following HLI were purified using magnetic bead separation. D) The number of satellite cells was increased in the ischemic limb (IL) compared to the nonischemic limb (NIL). E) Satellite cells in the ischemic limb had increased RAGE expression while satellite cells in the NIL has almost no expression. F) RAGE expression is enriched in satellite cells from the ischemic limb compared to the non-satellite cell fraction. (n=6, * indicates p<0.05)

We have previously shown that RAGE negatively impacts collateral vessel formation. (10) Therefore, we hypothesized that the satellite cells may have a counteracting effect influencing vascular regeneration through the production and secretion of a number of cytokines and growth factors. We used a multiplex immunoassay cytokine array to quantify cytokine expression in satellite cells from the ischemic and non-ischemic limbs of wild type mice. Both TNFα and MCP-1 protein levels were significantly increased in the ischemic limb (p<0.05), while IL-6 appeared to trend towards an increase but was non-significantly different (Figure 2 A-C). VEGF, however was not significantly different between the two limbs (data not shown). The levels of GM-CSF and IL-1β were below the detection range of our assay (1 pg/mL and 15 pg/mL respectively). We then compared the expression in satellite cells from the ischemic limbs of wild type and RAGE knockout mice. We found TNFα and MCP-1 were both significantly higher with p-valves of p=0.06 and 0.09 in the wild type cells compared to the RAGE knockout. IL-6 appeared to trend towards an increase but it was not significant and VEGF was also not different between the two cell groups (Figure 2 D-F).

**Figure 2:**
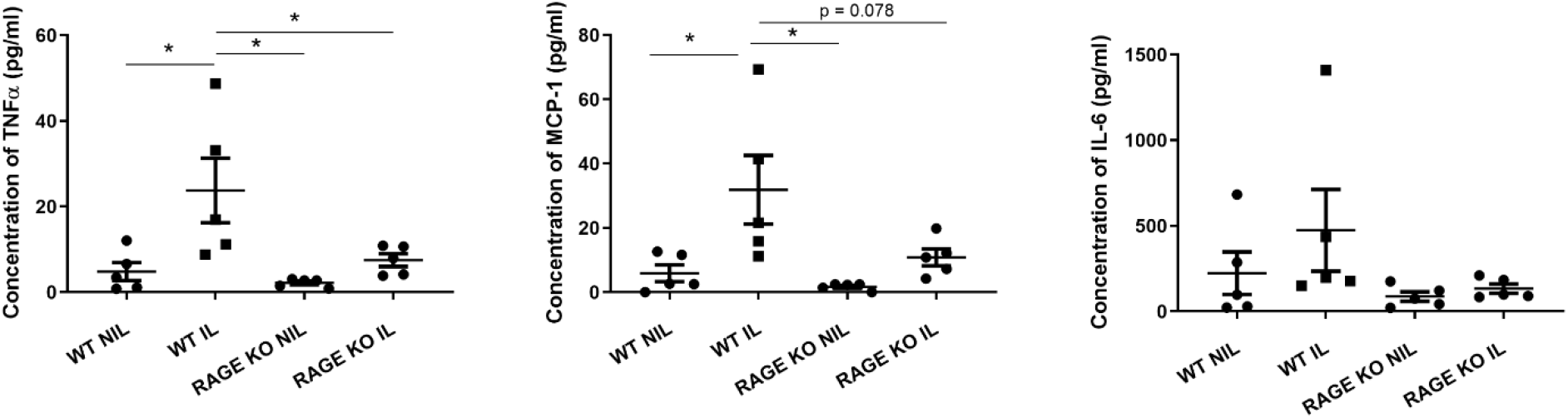
Wild Type but not RAGE Knockout Satellite Cells from the Ischemic Limb have Increased Cytokine Expression. Satellite cells were isolated using magnetic bead sorting from hind limb muscles at 7 days post HLI. Protein cytokine levels were quantified using a Uplex immunoassay (Meso Scale Diagnostics). A&B) Satellite cells from the ischemic limb (IL) had significantly great TNFα and MCP-1 protein production. (* indicates p<0.05) C) IL-6 followed similar trend but was non-significant. C) Satellite cells from the ischemic limb of RAGE knockout mice (RAGE KO IL) had significantly decreased TNFα expression compared to cells from wild type mice (WT IL) (p=0.06). E) RAGE knockout satellite cells also had significantly decreased MCP-1 expression compared to wild type cells. (p=0.09) F) IL-6 expression as not significantly different between the two cell types (p=0.19) (n=5)

We further examined the expression of these same factors in cultured wildtype and RAGE KO satellite cells in response to the RAGE ligand, HMGB-1 (Figure 3). Similar to the array data, stimulating wild type satellite cells (WT SCs + 250 nM HMGB1) increased expression of TNFα, MCP-1, and IL-6 compared to the stimulated RAGE knockout satellite cells (RAGE KO SCs + 250 nM HMGB1) (p<0.05).

**Figure 3:**
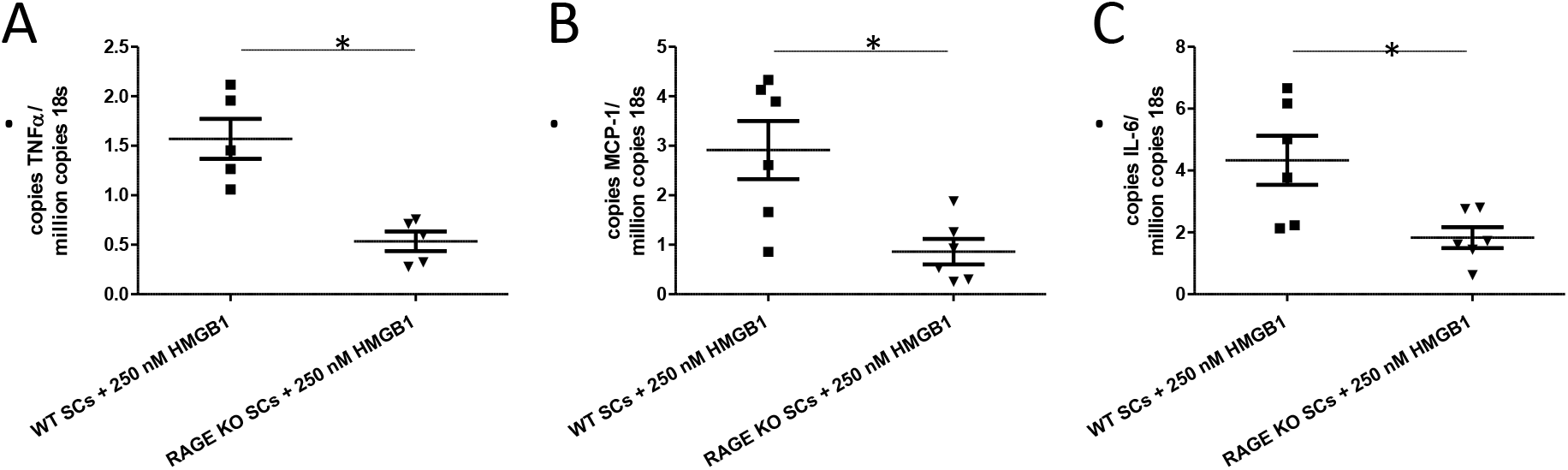
RAGE knockout satellite cells have decreased cytokine response to HMGB1 stimulation. Primary cultures of wild type or RAGE knockout satellite cells were stimulated with 250 nM HMGB1 and cytokine expression was quantified by PCR. A) RAGE knockout satellite cells had decreased TNFα expression compared to wild type satellite cells. B). MCP-1 expression was less in RAGE knockout satellite cells versus wild type satellite cells. C). RAGE knockout satellite cells also had significantly less IL-6 expression compared to wildtype cells. (n=6, * indicates p<0.05)

Next, we studied the effects of wild type versus RAGE KO satellite cells on collateral vessel formation in response to ischemia. Alginate encapsulated satellite cells were delivered near the proximal ligation point at the time of HLI surgery.(13, 14) The viability of the delivered cells was monitored by quantifying the luminescence emitted from luciferase expressing satellite cells. Figure 4 shows good viability for 10 days (approximately 50% of mean luminescence detected at day 0) with mean luminescence diminished by day 14 in most animals. The effects of vascular recovery were monitored via LDPI for perfusion and immunohistochemically staining for vessels. Figure 5 shows that animals that received wild type satellite cells had improved perfusion as measured by LDPI compared to animals that received empty capsules (p<0.05). The mice that received RAGE KO satellite cells did not show differences compared to the empty capsule treated group. In addition to LDPI, recovery was assessed by quantifying the lectin positive capillaries and small vessels as well as arteries and arterioles with smooth muscle actin via histological staining. Similar to the LDPI results, the group that received the wild type satellite cells had increased lectin positive vessels compared to the empty capsule control group with the group with RAGE knockout satellite cells not different from either of the other two groups (p=0.001) (Figure 6 A-D). Similarly, the smooth muscle actin staining showed an increase in arteries/arterioles in the mice that received wild type satellite cells compared to empty capsules (Figure 6 E-H).

**Figure 4:**
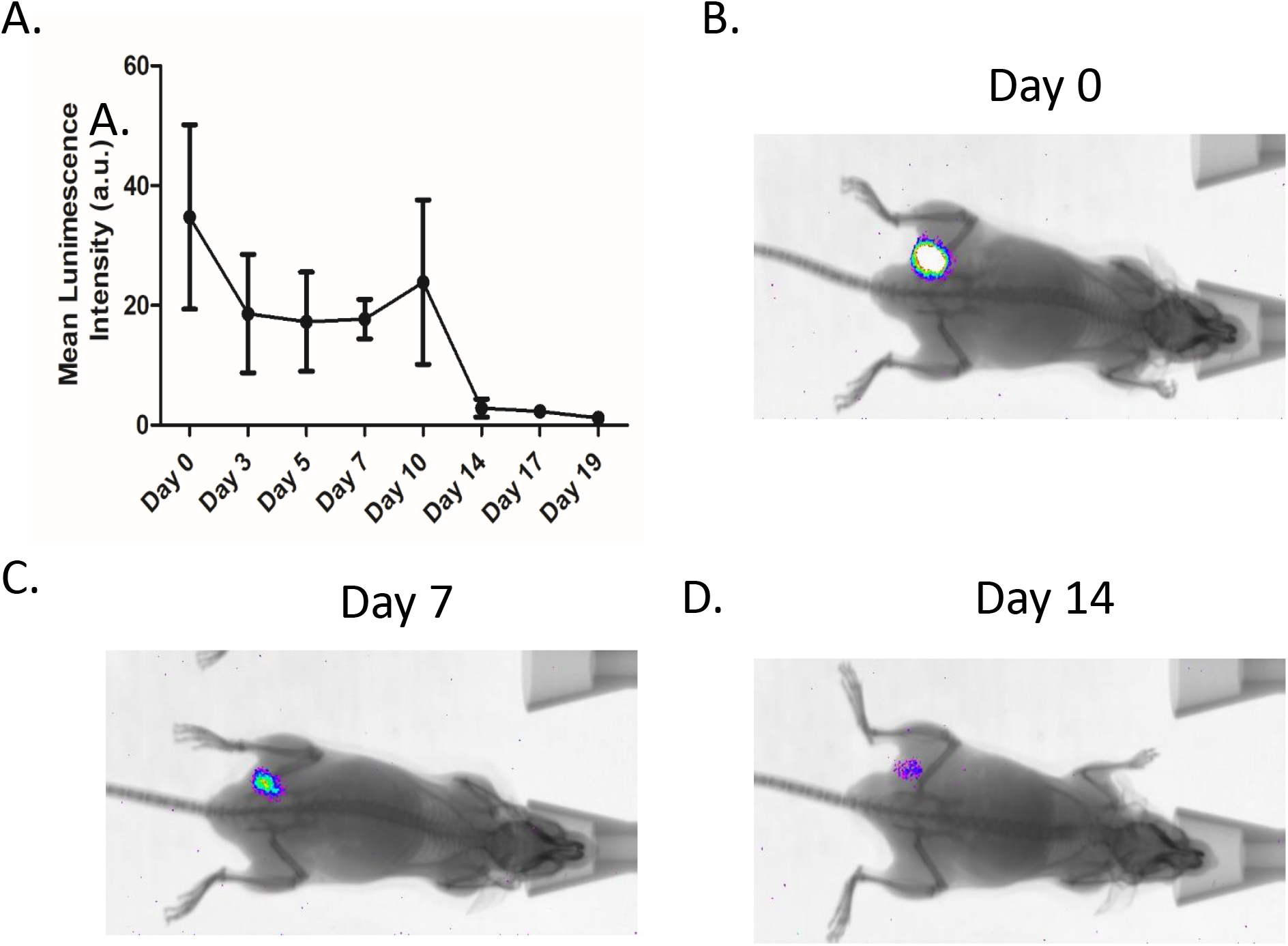
Implanted satellite cells are viable for >10 days. Luciferase expressing satellite cells were tracked in vivo using a whole animal in vivo imager (Bruker). A) Quantification of the luminescence showed good viability up to 10 days with some cells detected up to 19 days post implantation. B-D) Representative images at days 0, 7, and 14.

**Figure 5:**
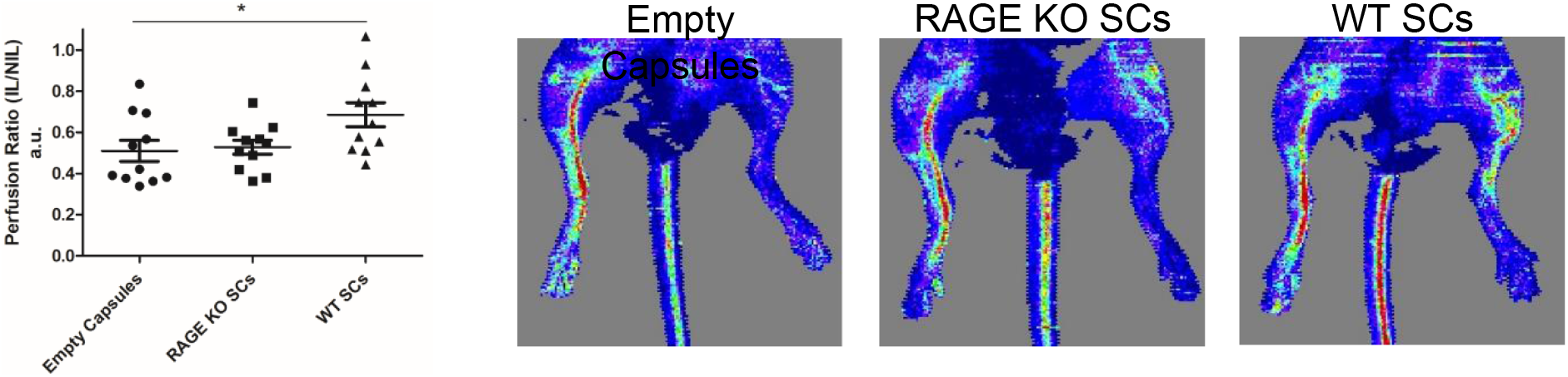
Delivery of Wild Type Satellite Cells Improves Perfusion in a Hind Limb Ischemia Model. Mice received either wildtype or RAGE knockout satellite cells in alginate capsules or empty capsules at the time of hind limb ischemia surgery. Perfusion recovery was measured via LASER Doppler perfusion imaging (LDPI) over time. Mice that received wildtype satellite cells (WT SCs) had improved perfusion over mice that received empty capsules or RAGE knockout satellite cells (RAGE KO SCs) at both 7 and 14 days post surgery. (n=11, * p<0.05 Empty Capsules vs WT SCs, Day 14 data shown)

**Figure 6:**
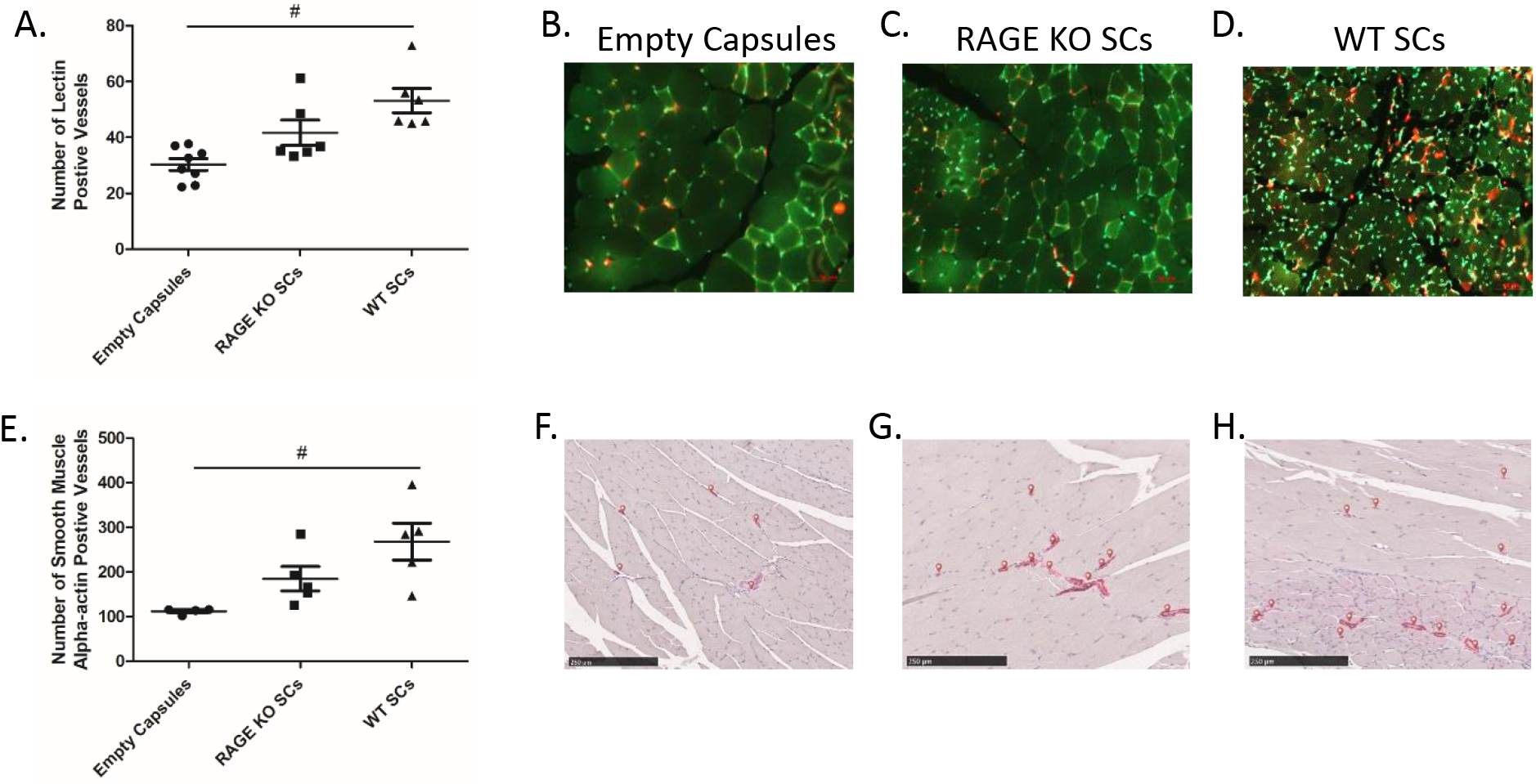
Delivery of wildtype satellite cells increases revascularization. Mice received either wild type or RAGE knockout satellite in alginate capsules or empty capsules at the time of hind limb ischemia surgery and vascular growth was assessed via immunohistochemistry at day 14 post surgery. A-D) Capillaries and small vessels were quantified using Lectin staining. Wildtype satellite cells resulted in increased small vessels over empty capsules. (n = 11, p = 0.001 for Empty Capsules vs WT SCs, Red = lectin) E-H) Arterioles and arteries were quantified by staining for smooth muscle alpha-actin. As with capillaries, animals that received wildtype satellite cells had more vessels than animals that received empty capsules. (n=5, p<0.05 for for Empty Capsules vs WT SCs, red arrows indicate vessels)

## Discussion

We have demonstrated the presence of a previously unappreciated role for RAGE-mediated satellite cell function in collateral vessel formation. In contrast to our previous studies showing that global RAGE deficiency led to improved collateral vessel function (10), the current work shows that RAGE stimulation of satellite cells exerts a positive and paradoxical effect on collateral vessel formation. We show that satellite cells release cytokines and growth factors previously shown to be enhance collateral vessel formation and that increased RAGE expression in the satellite cells extracted from ischemic tissue is associated with increased secretion of some of these factors. In addition to increased expression of cytokines by satellite cells in ischemic tissue, there is also an increase in the number of satellite cells in these tissues thus augmenting the potential impact of satellite cells on collateral vessel formation. Finally, our cell delivery studies suggest that satellite cells may prove to be a viable therapeutic strategy for ischemic tissues and this response is dependent on the presence of RAGE in the satellite cells.

Our finding that satellite cells are the primary source of RAGE in the ischemic limb is a novel finding as most work on the role of RAGE in vascular biology and ischemia have focused on RAGE expression by macrophages, endothelial cells, and smooth muscle cells.(11, 16–19) Yet, these findings do not exclude a variable role for RAGE signaling in vascular and inflammatory cells. However, our work is in agreement with work in muscle biology which show that while RAGE is expressed in skeletal muscle during development and in cultured myoblasts, healthy mature skeletal muscle has little basal RAGE expression.(17, 20) We also show in Figure 1 that satellite cells from the control limb had almost no RAGE expression while the cells in ischemic limb had robust expression and were increased in number. Thus, we sought to explore the role of satellite cell RAGE and its subsequent downstream signaling in our model of ischemic induced vascular growth.

We determined in addition to increasing in number, satellite cells from the ischemic hind limb muscles had increased TNFα and MCP-1 suggesting that ischemia stimulated satellite cells have a pro-inflammatory phenotype (Figure 2 A-C). We then compared the expression profiles of satellite cells from wild type and RAGE knockout mice and interestingly found that satellite cells from RAGE knockout satellite cells had decreased cytokine expression (Figure 2 D-F). We further explored these differences by stimulating cultured satellite cells with the RAGE ligand HMGB-1. Similar to the freshly isolated cells, in vitro stimulation of satellite cells produced more expression of cytokines TNFα, MCP-1, and IL-6 in wild type satellite cells versus RAGE knockout satellite cells (Figure 3). Taken together, the results in Figures 2 and 3 show that wild type satellite cells are responsive to stimuli with increased cytokine expression while RAGE knockout satellite cells are less responsive and express lower levels of cytokines overall. Thus, RAGE knockout satellite cells appear to have a less inflammatory phenotype when compared to wild type cells. This finding is contrary to the typical dogma that RAGE signaling leads to negative effects but is consistent with the concept that collateral vessel formation is driven by inflammatory responses. (18, 21–27) We speculate that stimulation of RAGE on satellite cells is an endogenous repair mechanism. Specifically the ligand HMGB1 which has been shown to be increased in the ischemic limb and may be responsible for RAGE signaling even in non-diabetic settings such as the hind limb ischemia model in this study.(10) HMGB1 is a transcription factor that is released from dead and damaged cells, thus the potential connection between released HMGB1 in the setting of ischemic tissue and stimulation of satellite cells makes teleological sense as a repair mechanism.

We hypothesized that the cytokine production by the satellite cells might play a role in regulating angiogenesis and arteriogenesis since it has been shown that increases in satellite cell number and capillary number are correlated in acute exercise induced muscle growth and muscle repair following injury.(28, 29) While previous studies have shown that RAGE knockout animals recover better in the hind limb ischemia model (10, 11), the results in Figures 2 and 3 show that satellite cells from the RAGE knockout mice have decreased production of several key cytokines and growth factors. Thus, we hypothesized that RAGE knockout satellite cells are less responsive to the ischemic insult, not capable of producing the same degree of cytokines and growth factors, and therefore less potent stimuli for vascular growth. To test this hypothesis, we delivered satellite cells to the ischemic limb at the time of surgery. The cells were encapsulated in alginate, which has been previously shown to improve the retention and survival of mesenchymal stem cells in both a hind limb ischemia and myocardial infarction model.(13, 14) Figure 4 shows that the satellite cells were viable for up to 10-14 days, which also correlates with the peak differences observed in LDPI analysis of perfusion and assessment of vessel number. This suggests the viability was sufficient to result in a positive effect on collateral vessel formation and function. In agreement with our hypothesis, the wild type satellite cells showed significant improvement over the empty capsules in several assessments of vascular growth while the animals with RAGE knockout satellite cells had no improvement or what appeared to be an intermediate effect that was not significantly different from either group (Figures 5 and 6).

While this finding of the wild type satellite cells having better outcomes than RAGE knockout cells may seem contrary to previous studies which have implicated RAGE as having negative effects (and thus positive effects when RAGE is knocked out), these studies focused on other cellular sources of RAGE.(6, 8–11, 30–32) This suggests that perhaps the negative effects of RAGE signaling are not due to satellite cells but other cells expressing RAGE. Importantly, our findings are consistent with multiple previous studies showing that RAGE signaling upregulates cytokine expression and secretion.(5, 27, 33) In Figures 2 and 3 we show that that RAGE knockout satellite cells produce less cytokines in both an in vivo and in vitro system. Specifically, others have shown that RAGE activates several mitogen-activated protein kinase (MAPK) signaling cascades, including p38, leading to NF-kB translocation, and signaling upregulation of multiple cellular responses including RAGE itself, IL-6 and TNFα (23, 33–37) We also showed that stimulation of wild type cells with a RAGE ligand leads to increases in cytokine expression including IL-6 and TNFα while the lack of RAGE produces a blunted response to the same stimuli (Figure 3).

While increased inflammation is typically viewed as a negative factor, it is known that some degree of inflammation and reactive oxygen species are required for proper collateral vessel formation.(38-44) Therefore, we purpose that satellite cells produce a physiological inflammatory response to ischemia and increased vascular growth and that RAGE is a critical mediator of this process. This suggests that in certain cell types, RAGE signaling might play a physiological rather than pathophysiological role in producing inflammatory signals. HMGB-1 mediated RAGE signaling in rat myoblasts has been shown to increase p38 phosphorylation which is required for myogenic differentiation and myogenesis. (17) In the C2C12 myoblast cell line, RAGE was shown to be required for differentiation through the p38-MAPK pathway to upregulate myogenesis and downregulate Pax7. (16) The MAPK pathway is upstream of myogenin, the transcription factor that marks terminal differentiation of the satellite cell into a myoblasts. Thus, satellite cells that lack RAGE fail to properly activate, proliferate, and differentiate into myoblasts, but instead have decreased differentiation and an increased number return to the quiescent state. (16, 17) This is consistent with dual function of RAGE-mediated inflammation in ischemic muscle injury leading to both muscle and vascular regeneration. It is important to note that in our study, the satellite cells were encapsulated in non-degradable alginate so while molecules are still able to diffuse to and from the cells, their lack of ability to differentiate and fuse to repair the muscle is not a factor in our results as only paracrine mechanisms are involved. These findings suggest that in addition to RAGE being required for proper renewal and differentiation of the satellite cell population, it is also required for production of cytokines and factors that drive vascular growth.

Our study demonstrates that in addition to their well known role in muscle repair, satellite cells play a very important role in regulating vascular growth. We found that ischemia both increases satellite cell number as well as changes their function to a more inflammatory phenotype. Interestingly, RAGE expression and signaling which is also increased in response to ischemia is an absolutely required mediator of this phenotypic change. Specifically, RAGE knockout satellite cells were not effective at increasing vascular regrowth. These findings corroborate with the literature that shows that the lack of RAGE changes the phenotype of the satellite cells and results in a more quiescent state. We believe that this phenotype change also results in decreased inflammatory signaling that impairs the vasculogenic potential of the satellite cells. Taken together it demonstrates that RAGE signaling is important for proper satellite cell function both in terms of muscle repair and coordinating the vascular response. Thus, while satellite cells appear to be a major source of RAGE in the ischemic hind limb, RAGE-mediated inflammatory responses appear to be required for their proper function. Furthermore, these studies raise the interesting possibility that administration of autologous satellite cells is a potential therapeutic strategy for the treatment of PAD.

## Abbreviations

HLI: Hind limb ischemia
HMGB1: high mobility group box 1
KO: knockout
LDPI: Laser Doppler perfusion imaging
PAD: peripheral artery disease
RAGE: Receptor of Advanced Glycation End Products
SCs: satellite cells
WT: wild type

## Highlights

Skeletal muscle satellite cells are novel source of RAGE in the ischemic hindlimb.

RAGE mediated cytokine production is critical for satellites cells’ role as a driver of vascular growth.

Satellite cells have the potential as a new therapeutic approach to treatment of peripheral artery disease.

